# miR-519a-3p, found to regulate cellular prion protein during Alzheimer’s disease pathogenesis, as a biomarker of asymptomatic stages

**DOI:** 10.1101/2023.12.15.569852

**Authors:** Dayaneth Jácome, Tiziana Cotrufo, Pol Andrés-Benito, Eulàlia Martí, Isidre Ferrer, José Antonio del Río, Rosalina Gavín

**Author notes:** Corresponding author: R. Gavín, PhD. **Senior co-authors: R. Gavín, PhD**, Department of Cell Biology, Physiology and Immunology, Faculty of Biology, University of Barcelona, Avinguda Diagonal 643, 08028 Barcelona, Spain, **J. A. del Río, Prof.**, MCN lab, Institute for Bioengineering of Catalonia (IBEC), Baldiri and Reixac 15-21, 08028 Barcelona, Spain.

## Abstract

MiRNAs induce post-transcriptional gene silencing by binding to the 3’-UTR of complementary messenger RNAs and causing either degradation or inhibition of translation.

The clinical relevance of miRNAs as biomarkers is growing due to their stability and detection in biofluids. In this sense, diagnosis at asymptomatic stages of Alzheimer’s disease (AD) remains a challenge since it can only be made at autopsy according to Braak NFT staging. Achieving the objective of detecting AD at early stages would allow possible therapies to be addressed before the onset of cognitive impairment.

Many studies have determined that the expression pattern of some miRNAs is deregulated in AD patients, but to date, none has been correlated with downregulated expression of cellular prion protein (PrP^C^) during disease progression. That is why, by means of cross studies of miRNAs up-regulated in AD with *in silico* identification of potential miRNAs-binding to 3’UTR of human *PRNP* gene, we selected miR-519a-3p for our study.

Other family members of miR-519 have been shown to bind to the 3’UTR region of *PRNP in vitro* and presumably degrade *PRNP* mRNA. In addition, up-regulation of some of them has been reported in various tissues from AD patients, including cerebrospinal fluid, plasma, and blood serum. In fact, miR-519d-3p is marked as a bridge regulator between mild cognitive impairment and severe AD. However, none of the studies address the prodromal stages of the disease or the expression profile of miR-519 in other neurodegenerative diseases that also may present dementia. Therefore, in this study we analyzed miR-519a-3p expression in cerebral samples of AD at different stages of evolution as well as other neurodegenerative diseases such as other tauopathies and synucleinopathies. Our results show the specific and early upregulation of miR-519a-3p starting from Braak stage I of AD, suggesting its potential use as a biomarker of preclinical stages of the disease.

## Introduction

### Alzheimer’s disease and its current diagnosis status

Alzheimer’s disease (AD) is far from having an early diagnosis, before mild or moderate cognitive deterioration appears and despite being the most common cause of dementia. This makes early therapeutic intervention very difficult to realize, which is currently restricted to 4 FDA-approved drugs unable to significantly reduce cognitive decline or improve global functioning.

As widely described, one of the neuropathological hallmarks of AD is extracellular accumulation of senile plaques, enriched in β-amyloid (Aβ) peptide. In addition, AD pathology courses like other tauopathies with intracellular aggregates of hyperphosphorylated tau protein into paired helical filaments (PHF), and then into neurofilament tangles (NFT). These are deposited progressively, starting in the entorhinal cortex and hippocampus and then spreading across other brain regions (Braak and Braak 1991; Iqbal, Braak et al. 1991; Braak, Braak et al. 1996; Avila 2000). In this sense, classical longitudinal studies of cognitive function and cerebrospinal fluid (CSF) analysis, as well as the switch from neuroimaging biomarkers used in dementia to AD-specific markers, have identified a significant preclinical phase of the disease that precedes onset of symptoms by about 10-20 years. This is characterized by early Aβ deposition in the precuneus and other cortical regions, followed sequentially by regional cortical hypometabolism, significant accumulation of tau pathology, and the onset of symptomatic cognitive impairment. Unlike Aβ, and as previously reported, the stage of tau pathology correlates well with the progression of cognitive decline (Serrano-Pozo, Frosch et al. 2011), largely following the Braak stages (I to VI) according to the regional distribution of NFT after *post-mortem* evaluation (Braak and Braak 1996).

In general terms, neuropathological diagnosis of AD can currently be made with reasonable certainty using CSF or positron emission tomography (PET) imaging biomarkers that allow estimation of cerebral Aβ and tau depositions (Tapiola, Alafuzoff et al. 2009; Lowe, Lundt et al. 2019). However, in some cases, cognitive impairment and behavioural changes are not apparent until severe AD dementia, preventing a reliable diagnosis. In addition, with the use of cognitive scales, such as the Mini-Mental Scale examination (MMSE), people in Braak stage I-II are asymptomatic and considered healthy, as shown by correlating the scores of the Braak staging and MMSE (Wischik, Harrington et al. 2014). Moreover, in clinical practice, there is no standardized criteria for diagnosis and, there may be serious variations between centres including heterogeneous methodologies as well as expensive and/or invasive ones (Hampel, O’Bryant et al. 2017; Hampel, Toschi et al. 2018; Dubois, Bombois et al. 2020). In this sense, the use of CSF carries the risks and inconveniences involved in a lumbar puncture procedure.

Thus, current trends are focused on the search for markers (some of them in the line of epigenetics; see below) in tissue extracted using non-invasive procedures (Zetterberg and Burnham 2019; Lee, Ugay et al. 2020). In this sense, the detection of peripheral biomarkers in samples such as blood is emerging as an advance in terms of the correlation of classical biomarkers in CSF and PET images (de Rojas, Romero et al. 2018; Risacher, Fandos et al. 2019) (collected at (Leuzy, Mattsson-Carlgren et al. 2022)). However, general population screening continues to be a long-term vision (Teunissen, Verberk et al. 2022).

Among epigenetic signatures in AD, a large number of studies show the dysregulation of various microRNAs (miRNAs) in the disease. Importantly, the potential of miRNAs as diagnostic tools has been widely reported to show a regular pattern among various body fluids (Hanna, Hossain et al. 2019; Cui and Cui 2020) and a marked correlation between the expression levels in plasma and brain parenchyma (Kos, Puppala et al. 2022). In this regard, miR-519a-3p has already been proposed as a biomarker of AD in patients showing mild cognitive impairment (MCI) (Jia and Liu 2016; Lusardi, Phillips et al. 2017; Zhao, Zhang et al. 2019; Tao, Han et al. 2020) and it has been found in various fluids such as CSF, blood, plasma, and serum. However, its potential as a biomarker of asymptomatic stages (according to Braak stages I and II) has never been analyzed before.

### Epigenetic signature of cellular prion protein as possible blood biomarker for AD

The relationship between cellular prion protein (PrP^C^) and AD is widely reported. PrP^C^ was initially known to be the causative agent of prionopathies when it undergoes changes in its normal folding (Prusiner 1982). However, its physiological role is still a subject of debate and some studies show its interaction/participation with proteins or signalling pathways involved in AD (see some examples in (Parkin, Watt et al. 2007; Griffiths, Whitehouse et al. 2012; Nieznanska, Boyko et al. 2021)). Even so, the role of PrP^C^ in the disease continues without being resolved, since some studies grant it a pathogenic role (Lauren, Gimbel et al. 2009) while others support a neuroprotective function (e.g. (Westergard, Christensen et al. 2007) reviewed in (Gavin, Lidon et al. 2020)).

PrP^C^ is mainly expressed by neurons and glial cells in the adult central nervous system (CNS) (Moser, Colello et al. 1995; Ford, Burton et al. 2002; Moleres and Velayos 2005) and, in humans it is encoded in a single gene, *PRNP*, whose sequence is highly conserved in vertebrates (Nicolas, Gavin et al. 2009). Relevantly, the PrP^C^ expression profile varies during development (Moser, Colello et al. 1995; Miele, Alejo Blanco et al. 2003; Miranda, Ramos-Ibeas et al. 2013) with ageing in adults (Whitehouse, Jackson et al. 2010) and during AD progression, showing higher levels of expression in asymptomatic stages of the disease (McNeill 2004; Rezaie, Pontikis et al. 2005; Vergara, Ordonez-Gutierrez et al. 2015) and a progressive reduction in advanced stages.

To date there is no known mechanism involved in the reduction of PrP^C^ levels in the progression of AD, although abundant miRNA binding sites in the *PRNP* 3’UTR are known to be responsible (Pease, Scheckel et al. 2019). The aforementioned study showed the binding of some miR-519 family members (b-3p and d-3p) to the *PRNP* 3’UTR.

Therefore, in this study, we first cross-checked data from miRWalk, miRanda, RNA22, and Targetscan databases to select putative miRNAs responsible for the reduction of PrP^C^ during AD progression, pointing up miR-519a-3p as a candidate. Then, we demonstrated *in vitro* functional reduction of PrP^C^ by miR-519a-3p using mimic technology. Secondly, we analysed miR-519a-3p expression in frontal cortex samples of AD (from Braak I to VI) and other neurodegenerative disease (NDD) such as non-AD tauopathies and synucleopathies. Thus, we revealed early and exclusive expression of the miR-519a-3p in AD as a useful tool to generate an AD-specific signature in asymptomatic stages of the disease.

## Materials and Methods

### miR-519a-3p selection for the study

*In silico* prediction of potential miRNAs with *PRNP* 3’UTR as target was made crossing data from the miRWalk, miRanda, RNA22, and Targetscan databases. After that, candidates were analyzed for reported overexpression in AD (Lau, Bossers et al. 2013). Finally, miR-519a-3p (AAAGUGCAUCCUUUUAGAGUGU) was selected as the main candidate taking into account that it has already been found in various fluids such as CSF, blood, plasma, and serum in patients with mild cognitive impairment (MCI) and AD (Jia and Liu 2016; Lusardi, Phillips et al. 2017; Zhao, Zhang et al. 2019; Tao, Han et al. 2020).

### Cell culture and transfection

CCF-STTG1 human grade IV astrocytoma cells (CRL-1718, American Type Culture Collection (ATCC)) were used to analyse the levels of PrP^C^ *in vitro* after has-miR-519a-3p Mimic (Qiagen) transfection. Cells were maintained in Roswell Park Memorial Institute (RPMI) 1640 medium supplemented with 10% fetal bovine serum (FBS), and 1% penicillin/streptomycin (Thermo Fisher Scientific, MA, USA) in 75 cm^2^ culture bottles (Nunc, Denmark) in a 5% CO_2_ atmosphere at 37ºC. One week before transfection, cells were cultured in the same medium, on poly-D-lysine (Sigma-Aldrich, Darmstadt, Germany) coated 6-well plates (Nunc, Denmark).

HEK293 human embryonic kidney cells (ATCC CRL-1573, American Type Culture Collection, Rockville, Md., USA) were used to perform *in vitro* miR-519a-3p target analysis on 3’UTR-*PRNP* reporter construct (vector pEZX-MT06, Genecopoeia). Cells were maintained in Advanced Dulbecco’s modified Eagle’s medium (AdDMEM) supplemented with 10% fetal bovine serum (FBS), 0.5% glutamine, and 1% penicillin/streptomycin (Thermo Fisher Scientific, MA, USA) in 75 cm^2^ culture bottles (Nunc, Denmark) in a 5% CO_2_ atmosphere at 37ºC. One day before cotransfection, cells were cultured in the same medium, on poly-D-lysine (Sigma-Aldrich, Darmstadt, Germany) coated 24-well plates (Nunc, Denmark).

Transfections were performed using Lipofectamine 2000 (Invitrogen-Thermo Fisher Scientific) according to the manufacturer’s instructions.

### Human samples

The brains of non-neurodegenerative controls (nND) and patients with AD, other tauopathies non-AD, and synucleinopathies were obtained after death and were immediately prepared for morphological and biochemical studies.

A total of 106 frontal cortex (area 8) postmortem samples were obtained from Clinic-IDIBAPS, HUB-ICO-IDIBELL and NAVARRABIOMED Biobanks following the guidelines of Spanish legislation on this matter and approval of the local ethics committees. Total cases comprised nND (n = 14), AD (n = 61), non-AD tauopathies (n = 10), and synucleinopathies (n = 21).

Following neuropathological examination, AD cases were categorized according to neurofibrillary tangles (stages I to VI) and Aβ deposition (stages A, B and C) (Braak and Braak 1996; Braak, Braak et al. 1998). Non-AD tauopathies comprise corticobasal degeneration (CBD), (n = 4), glial globular tauopathy (GGT), (n = 3), and Pick’s disease (PiD) (n = 3). Synucleinopathies comprised Parkinson’s disease (PD, n = 21), 3 of them with associated dementia (PDD, n = 3). Neuropathological staging of these cases was based on the classification of Braak, from 1 to 6 (Braak, Del Tredici et al. 2003). Finally, nND cases did not show neurological or metabolic disease, and the neuropathological examination, carried out in similar regions and with the same methods as in pathological cases of this study, did not show lesions. In particular, no amyloid, tau or α-synuclein deposits were seen in the regions examined. A summary of basic patient data is presented in Supplementary Table 1.

### RT-qPCR

Total RNA from human hippocampal/frontal cortex samples and cultured cells was extracted with mirVana’s isolation kit (Ambion, TX, USA) following the manufacturer’s instructions. Total purified RNAs were used to generate the corresponding cDNAs, which served as PCR templates for qPCR assays. Before that, total RNA was quantified by qubit and analyzed for RNA integrity (RIN) in genomics service of Scientific and Technological services of the UB (CCiTUB). Retro transcription (RT) was performed using the miRCURY LNA RT kit (Qiagen, Germany).

Quantitative PCR (qPCR) for miR-519a-3p was performed in triplicate using miRCURY LNA SYBR Green PCR kit (Qiagen) and hsa-miR-519a-3p, LNA™ PCR primer set (Qiagen). PCR amplification and detection were performed with the Applied Biosystems StepOnePlus Real-Time detector. The following thermal profile was applied: 1 cycle at 95 °C for 2 minutes, 40 cycles at 95 °C for 10 seconds, and 56 °C for 60 seconds. Melting curve analysis was performed at 95 °C for 15 seconds, 60 °C for 60 seconds, and 95 °C for 15 seconds. miRNA levels were calculated using the StepOne™ software v2.3, following the 2^-*ΔΔC*T^ method of Applied Biosystems (Livak and Schmittgen 2001). Samples were normalized for the relative expression of the housekeeping miR103a-3p (hsa-miR-103a-3p, LNA™ PCR primer set, Qiagen), which showed no variability between analyzed groups.

RT-qPCR for *PRNP* mRNA was performed in triplicate using the following primers: (5’-agtcgttgccaaaatggatca-3’) and (5’-aaaaaccaacctcaagcatgtgg-3’) (Bribian, Fontana et al. 2012). PCR amplification and detection were performed with the Applied Biosystems StepOnePlus Real-Time detector, using 2x SYBR GREEN Master Mix (Roche Diagnostic, Switzerland) as reagent, following the manufacturer’s instructions. The reaction profile was denaturation-activation cycle (95ºC for 10 minutes) followed by 40 cycles of denaturation-annealing-extension (95ºC for 15 seconds, 60ºC for 60 seconds), and finally the melting curve analysis performed at 95 °C for 15 seconds, 60 °C for 60 seconds, and 95 °C for 15 seconds. mRNA levels were calculated using the StepOne™ software v2.3, following the 2^-*ΔΔC*T^ method of Applied Biosystems (Livak and Schmittgen 2001). The results were normalized for the expression levels of the housekeeping human *GAPDH* genes (5’-aggtcggtgtgaacggatttg-3’) and (5’-tgtagaccatgtagttgaggtca-3’), which were quantified simultaneously with the target gene (Carulla, Bribian et al. 2011).

### Western blotting techniques

Soluble extracts from CCF-STTG1 cultured cells were processed for western blot. Cells were washed in 0.1 M phosphate-buffered saline (PBS) before collection in sample buffer, Laemmli (Sigma-Aldrich, Darmstadt, Germany). Then, homogenates were boiled at 95ºC for 10 minutes, followed by 12% SDS-PAGE electrophoresis, and they were then electrotransferred to nitrocellulose membranes for 1 hour at 4ºC. Membranes were blocked with 5% non-fat milk in 0.1 M Tris-buffered saline (pH 7.4) for 1 hour and incubated overnight in 0.5% blocking solution containing primary antibodies. Monoclonal anti-PrP 6H4 (Prionics, Zurich, Switzerland) was used to determine PrP^C^ levels and monoclonal anti-α-actin (Merck Millipore) was used for standardization. After incubation with peroxidase-tagged secondary antibodies (1:5000 diluted), membranes were revealed with the ECL-plus chemiluminescence western blot kit (Amersham-GE Healthcare, UK).

For western blot quantification, developed films were scanned at 2,400 x 2,400 dpi (i800 MICROTEK high quality film scanner) and the densitometric analysis was performed using Fiji™ software (National Institutes of Health, USA).

### miR-519a-3p target validation with dual-luciferase reporter assay

To determine targeting of miR-519a-3p to *PRNP* 3’UTR miRNA sequence, 4.5 x 10^4^ HEK293 was seeded 24 hours before transfection into 24-well plates. Cells were cotransfected with human *PRNP* 3’UTR miRNA Target Clone (pEZX-3’UTR-PRNP vector from Genecopoeia) and hsa-miR-519a-3p miRCURY LNA miRNA Mimic or its related Negative Control miRCURY LNA miRNA Mimic (Qiagen). Cell lysates were prepared 24 hours later in passive lysis buffer following the manufacturer’s instructions, and luciferase reported activity was measured with an Infinite M200 Pro Microplate Reader luminometer using the Dual-Luciferase assay system (Promega). Background luminosity was subtracted from un-transfected cells and luciferase readings were normalized by dividing the reported gene firefly luciferase by the reference Renilla luciferase activity. Each construct was transfected in triplicate in three independent experiments.

### Statistical processing

Data analysis was performed using Prism 8.0 (GraphPad Software, CA, USA). Data were obtained using Student t-test. Differences with *p* < 0.05, *p* < 0.01, *p* < 0.001, and *p* < 0.0001 were considered significant. All data are expressed as mean ± standard error of the mean (SEM).

## Results

### miR-519a-3p overlaps PRNP 3’UTR in vitro and promotes changes in PrP^C^

After identifying of miR-519a-3p as a candidate to induce downregulation in PrP^C^ during AD, we explored its functional activity on PrP^C^ expression. To do this, we performed two parallel *in vitro* experiments using mimic technology. First, we analysed target of miR-519a-3p to *PRNP* 3’UTR using the pEZX-MT06-3’UTR-*PRNP* vector for Firefly luciferase expression under 3’UTR-*PRNP* control and Renilla luciferase for standardization. 24 hours after HEK293 co-transfection with miR-519a-3p mimic or its corresponding negative control and 3’UTR-*PRNP* construction, a dual-luciferase assay was performed. Results showed a significant decrease of 21.12% ± 1.40 of Firefly bioluminescence when miR-519a-3p mimic was expressed, confirming predicted 3’UTR-*PRNP* site as target for miRNA (Figure 1A).

**Figure 1.**
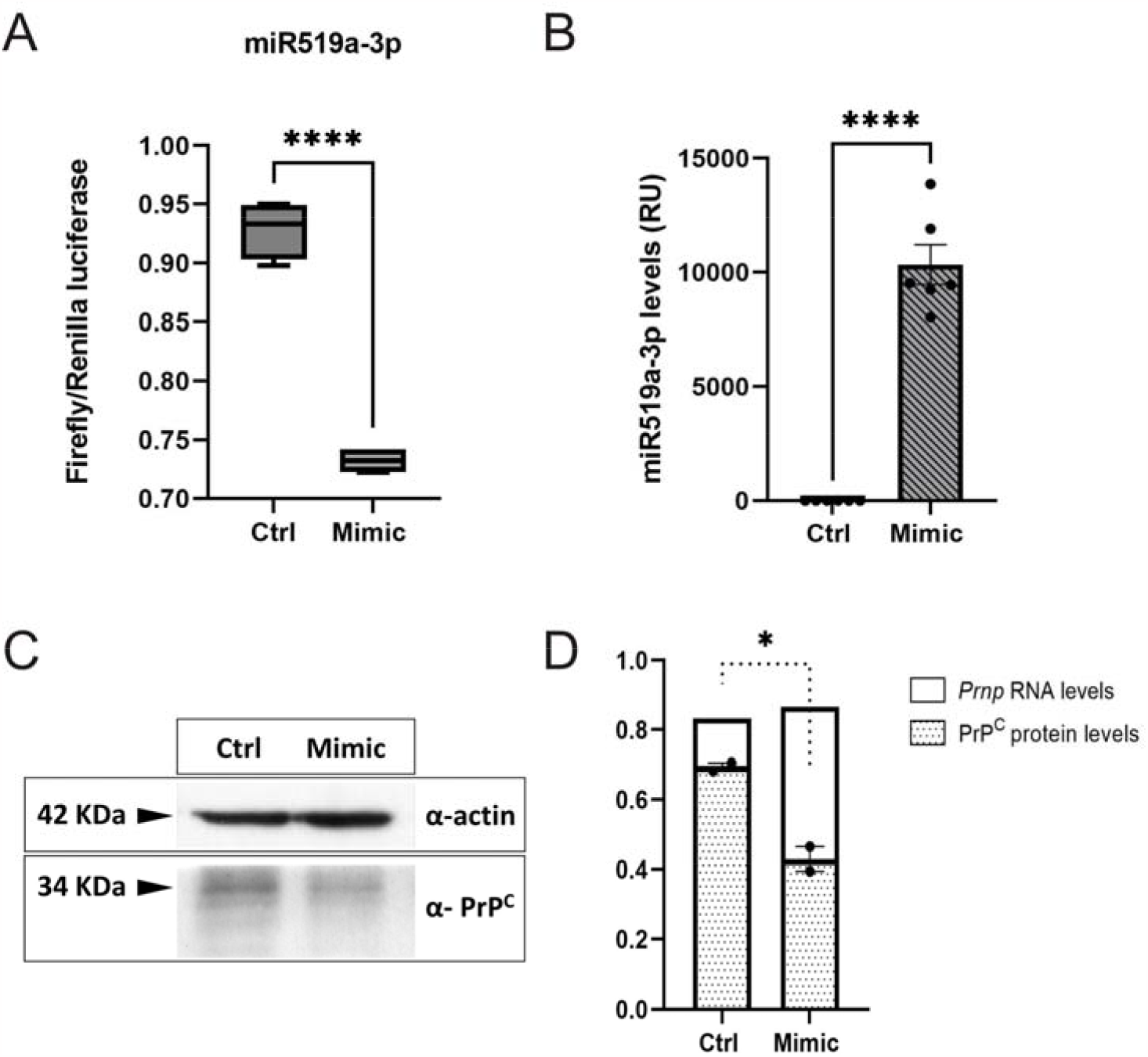
Binding of miR-519a-3p mimic to the *PRNP* 3’UTR and decrease in PrP^C^ levels. Dual-luciferase reported assay showed the binding of miR-519a-3p to the *PRNP* 3’UTR after co-transfection of HEK293 cells with mimic and pEZX-3’UTR-*PRNP* vector by reduction of Firefly luciferase activity **(A)**. Human CCF-STTG1 cell line shows overexpression of miR-519a-3p 48h after transfection **(B)** and it is associated with a decrease of around 25% in PrP^C^ protein levels **(C, D)** while mRNA levels remain unchanged **(D)**. PrP^C^, *PRNP* mRNA and miR-519a-3p were analyzed with Western blot and RT-qPCR respectively. Differences between groups were considered statistically significant at ^****^*p* < 0.0001 and ^*^*p* < 0.05 (Student’s t-test). RU: Relative units.

It is reported that miR-519a-3p only works on human genome, so the second approach was developed on *CCF-STTG1* cell line, a human astrocytoma line which presents high expression of PrP^C^. Human cell line was transfected with miR-519a-3p mimic and with the negative control in parallel to measure later levels of both PrP^C^ protein and mRNA *PRNP*. Then, both RNA and protein were extracted at 48h after mimic transfection. Firstly, our results showed a significant increase in expression of miR-519a-3p by RT-qPCR (Figure 1B). And higher levels of miR-519a-3p were associated with a significant decrease of PrP^C^ with western blot analysis (Figure 1C), which corresponded to 38.02% ± 3.74 while mRNA *PRNP* levels, analyzed with RT-qPCR, remained unchanged (Figure 1D), suggesting translational repression at this time point.

### Significant changes in miR-519a-3p levels at subclinical stages of AD

In order to analyze miR-519a-3p levels in the AD sample collection under study we performed an RT-qPCR. Then, data were classified into 3 groups according to Braak NFT stage of patients, without taking into account stages of Aβ deposition (Supplementary Table 1). Importantly, quantitative results showed an increase in all three groups when compared to nND samples: AD(I/II) (1.883 ± 0.258; ^**^ *p* = 0.0012), AD(III/IV) (2.484 ± 0.303; ^****^ *p* = <0.0001) and

AD(V/VI) (2.768 ± 0.554; ^**^ *p* = 0.0012) (Figure 2). In addition, although there was a tendency to increase with the progression of the disease, we did not find *statistically significant differences between the 3 groups of affected people*.

**Figure 2.**
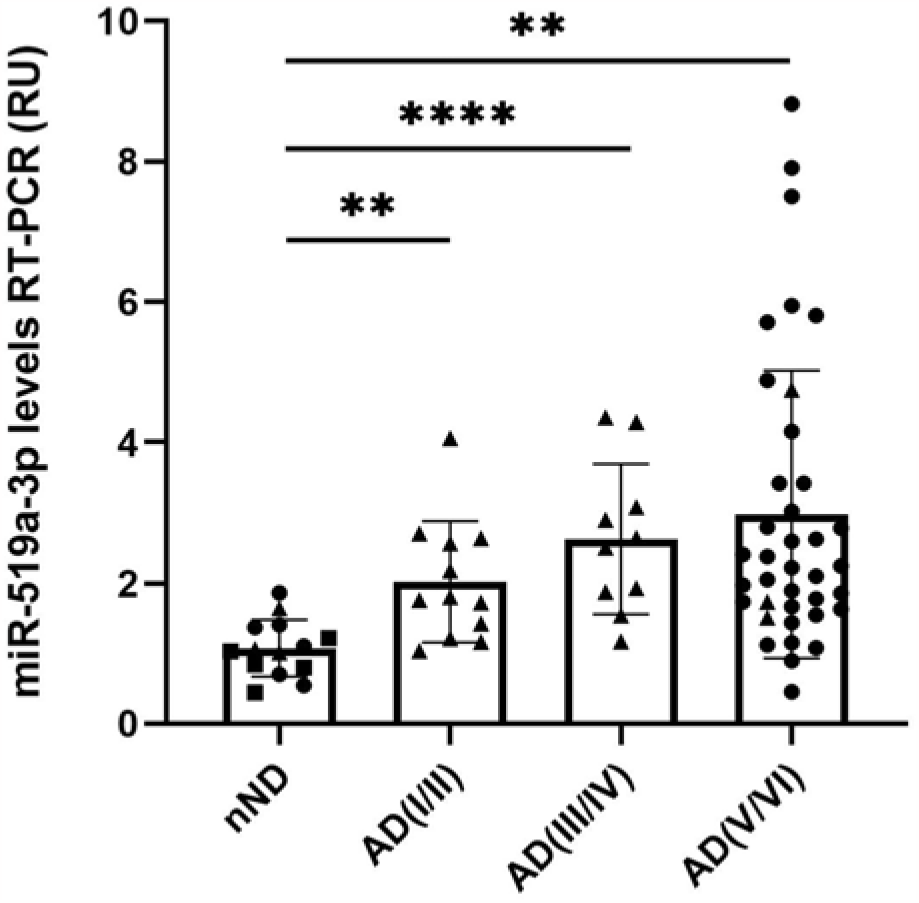
miR-519a-3p expression in frontal cortex samples with AD grouped according to NFT Braak staging (AD(I/II), AD(III/IV), and AD(V/VI)) and compared to nND samples. The bars represent the mean of miRNA levels ± SEM between each group analyzed. Each dot represents a case and each shape the biobank of origin according to circle: BioBANC Clinic-IDIBAPS; triangle: HUB-ICO-IDIBELL Biobank and square: NAVARRABIOMED Biobanc. Differences between nND samples and the three AD groups were considered statistically significant at ^****^*p* < 0.0001 and ^**^*p* < 0.01 (Student’s t-test). RU: Relative units.

### Specific pattern of miR-519a-3p expression in AD when compared with other NDD

To further explore specific dysregulation of miR-519a-3p in AD, we analysed the miRNA levels in additional samples from other NDD that occur with accumulation of fibrillar proteins as in AD and in some cases with dementia as well, first, in several advanced non-AD tauopathies including a few representations of CBD, GGT and PiD and second, in various synucleopathies including PD and some cases of PDD and classified into two groups according to staging of brain pathology related to sporadic PD. Thus, these groups are represented as PD(3-4) and PD(5-6), respectively (Supplementary Table 1).

Quantification of miR-519a-3p levels in advanced non-AD tauopathies compared to nND resulted in a non-significant increase of 1.282 ± 0.1999; *p* = 0.1419 (Figure 3A). Similarly, we found no significant differences in either of the two groups of PD analyzed, despite the fact that some of them presented dementia (Figure 3B). In this sense, fold change of PD(3-4) compared to nND was 1.141 ± 0.2050; *p* = 0.4673 while fold change of PD(5-6) compared to nND was 1.37 ± 0.2030; *p* = 0.0705.

**Figure 3.**
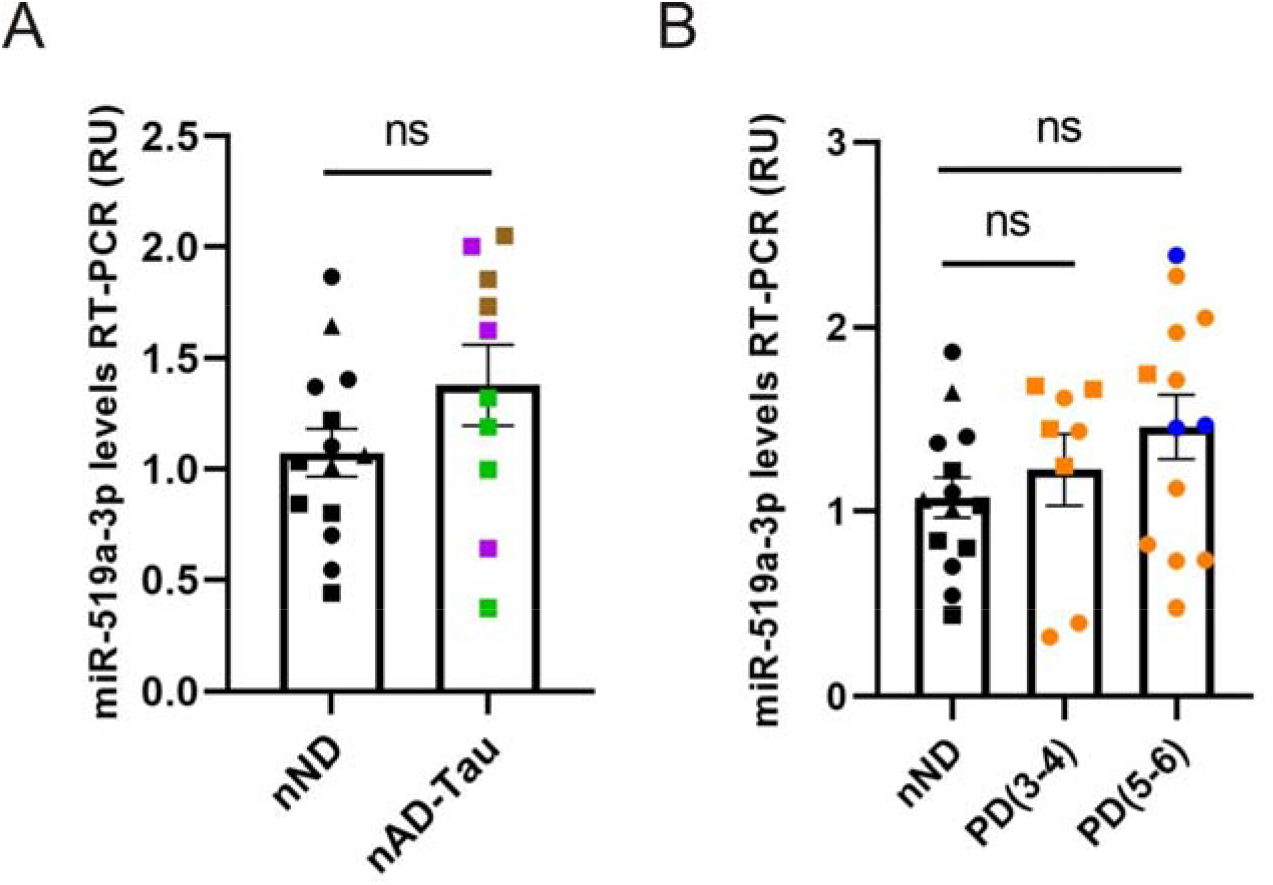
miR-519a-3p expression in non-AD tauopathies (and-Tau) (A) and Parkinson’s disease classified in PD(3-4) and PD(5-6) (B) compared to nND samples. Bars represent the mean of miRNA levels ± SEM between groups analyzed. Each point represents a case while each shape corresponds to the Biobank of origin according to circle: BioBANC Clinic-IDIBAPS; triangle: HUB-ICO-IDIBELL Biobank and square: NAVARRABIOMED Biobanc. In addition, the colour codes represent the neuropathological diagnoses of the samples according to nND: Black, PD: Blue, PDD: Orange, CBD: Brown, GGT: Violet, PiD: Green. Differences between nND samples with nAD-Tau, as with PD(3-4) and PD(5-6), were not considered statistically significant (Student’s t-test). RU: Relative units.

Finally, we compared the AD cases distributed in groups AD(III/IV) and AD(V/VI) with the other NDD groups, non-AD tauopathies, PD(3-4), and PD(5-6), respectively. As observed in Figure 4, the two AD groups showed much higher and significant values of miR-519a-3p than the other NDD groups analyzed. Specifically, the fold increase in the AD(III/IV) group was 2.176 ± 0.4074; *p* = 0.0027 when compared to PD(3-4) and 1.936 ± 0.3749; *p* = 0.0029 when compared to non-AD tauopathies. In addition, the increase in the AD(V/VI) group was 2.041 ± 0.5795; *p* = 0.0117 when compared to PD(5-6), and 2.158 ± 0.6578; *p* = 0.0192 when compared to non-AD tauopathies. Therefore, in all cases the results are alike when comparing AD with the nND controls.

**Figure 4.**
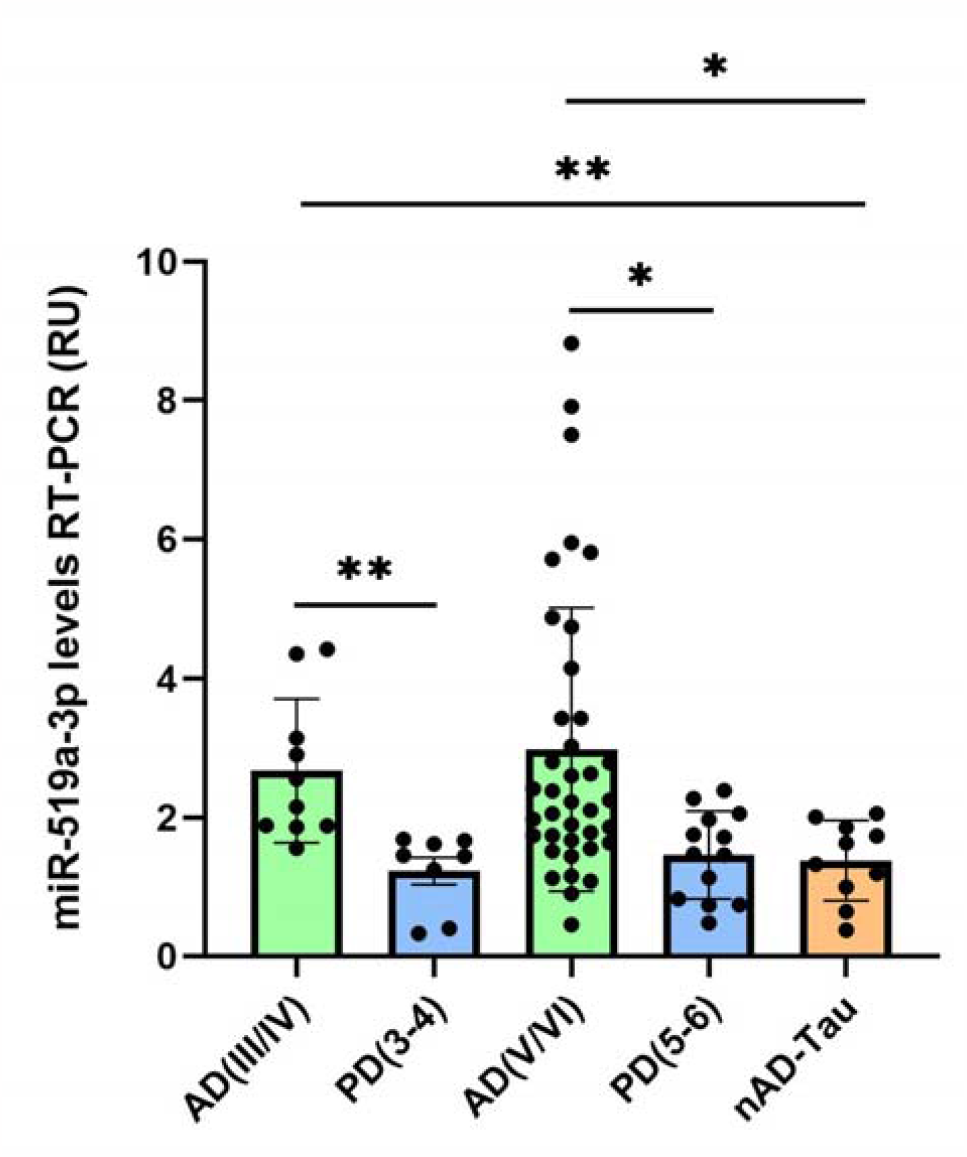
miR-519a-3p expression in AD samples distributed in AD(III/IV) and AD(V/VI) groups and compared to nAD-Tau, PD(3-4), and PD(5-6) groups respectively. Bars represent the mean of miRNA levels ± SEM between each group analyzed. Each point represents one case. Differences between AD, PD and and-Tau samples were considered statistically significant at \*\**p* < 0.01 and \**p* < 0.05 (Student’s t-test). RU: Relative units.

Taken together, these results suggest that elevated levels of miR-519a-3p could be considered in the search for a specific AD biomarker.

## Discussion

In the present study, we investigated two main issues concerning the role of miR-519a-3p in AD. On the one hand, there is its possible involvement in reducing PrP^C^ expression levels in the evolution of the disease (Vergara, Ordonez-Gutierrez et al. 2015; Lidon, Llao-Hierro et al. 2021). In fact, the role of PrP^C^ in AD and other NDDs is still a matter of debate, as some argue for its neuroprotective function while others point to it as an enhancer of the neurotoxicity associated with NDD (discussed in (Gavin, Lidon et al. 2020)). That is why it is important to identify the factors that modulate its expression levels, causing the protein to show increased levels in the brain parenchyma of patients in asymptomatic stages of AD and decreasing levels as the disease progresses.

The second issue to investigate is the potential use of this miRNA as a biomarker for the detection of AD in its asymptomatic stages. Up-regulation of some family members of miR-519 has been widely reported in different tissues from AD patients, e.g. FC (Lau, Bossers et al. 2013), CSF (Lusardi, Phillips et al. 2017) and in blood (plasma or serum) (Jia and Liu 2016; Zhao, Zhang et al. 2019; Tao, Han et al. 2020). In fact, the latter points to miR-519d-3p as a bridge regulator between MCI and AD. However, scant attention has been paid to analyzing upregulation of any member of the miR-519 family in asymptomatic people who may have initiated the disease silently (with Braak stage I-II correlation) (Wischik, Harrington et al. 2014). In our study, *in silico* prediction of potential miRNAs with *PRNP* 3’UTR as target pointed to miR-519a-3p (Lau, Bossers et al. 2013) among others as a candidate up-regulated in AD. Importantly, binding of other miR-519 family members (b-3p and d-3p) to *PRNP* 3’UTR and the consequent mRNA degradation has already been demonstrated *in vitro* (Pease, Scheckel et al. 2019). In this sense, it is further assumed that mature miRNAs belonging to the same family share the same target genes according to conserved regions in their sequences (Cai, Yu et al. 2009). Specifically, miR-519a-3p and miR-519b-3p belong to the same subfamily, so they share exactly the same seed sites, while miR-519d-3p is part of the AGUGC family like both of them (Wang 2009). Thus, our results using mimic technology corroborate the control of PrP^C^ expression by this additional miR-519 family member, first, determining miR-519a-3p target validation with dual-luciferase reporter assay, and secondly, downregulating PrP^C^ protein levels after miR-519a-3p mimic overexpression. However, in contrast to previous data, our experiment elicited translational repression and not mRNA degradation, since *PRNP* mRNA levels did not change. Perhaps the time point at which we measured PrP^C^ protein and *PRNP* mRNA levels was decisive in this regard because we analyzed the samples 48h after mimic transfection while they did so at 72h (Pease, Scheckel et al. 2019).

In addition, after analyzing the miR-519a-3p levels in the 61 frontal cortex samples of AD, we corroborated the significant up-regulation in Braak III-IV compared to non-AD samples. This is why these results support the possible use of miR-519a-3p in the detection of patients presenting MCI as suggested in (Tao, Han et al. 2020). However, our results also show that this miRNA is significantly up-regulated in apparently asymptomatic Braak I-II samples, as identified at the time of the post-mortem neuropathological examination. In fact, elevated levels of miR-519a-3p have been previously reported in prefrontal cortex of late-onset AD patients from Braak I (Lau, Bossers et al. 2013), although the focus of the aforementioned study was on the identification of miRNAs directly implicated in pathogenesis of the disease, and no attention was paid to possible biomarkers. This finding, together with its already reported detection in blood of AD patients (Zhao, Zhang et al. 2019; Tao, Han et al. 2020), makes miR-519a-3p a putative diagnostic tool for the disease.

In this sense, there currently exist several blood-based biomarkers that correlate well with the classical diagnosis of the disease by means of CSF analysis or imaging tests. Specifically, MRI for the detection of brain atrophy and PET with radiotracers for the detection of Aβ or phosphotau deposits (Cullen, Leuzy et al. 2021; Leuzy, Cullen et al. 2021; Ossenkoppele, Reimand et al. 2021; Zetterberg and Bendlin 2021; Teunissen, Verberk et al. 2022). However, these emerging biomarkers cannot detect pre-clinical disease since differential levels of them are detectable only when the degenerative process starts showing signs of dementia (Ferrer 2023).

Several studies show the potential of miRNAs as diagnostic tools for their stability and abundance in circulatory fluids (reviewed in (Hanna, Hossain et al. 2019). The ‘golden’ indicator of RNA degradation is the ‘RNA integrity number’ (RIN) that cannot be applied to miRNAs which are extremely stable against nucleases (Jung, Schaefer et al. 2010). In this sense, our results have shown the non-association between RIN and miR-519a-3p level that only depends on AD evolution (Supplementary Figure 1). But more importantly, miRNAs show a regular pattern of expression among various body fluids including blood-based fluids, which facilitates their extraction with non-invasive methods for diagnostic use (Cui and Cui 2020). In addition, miRNAs can cross the blood brain barrier (BBB) (Beylerli, Gareev et al. 2022), and correlation between miRNA expression levels in plasma and brain parenchyma has even been reported (Kos, Puppala et al. 2022).

Although other neurodegenerative diseases such as PD and tauopathies may present with dementia like AD, most of them show diagnoses well-differentiated from AD. This is not always the case, as hippocampal atrophy on magnetic resonance imaging (MRI) is an early characteristic of AD that may also occur in other dementias, such as FTLD (van de Pol, Hensel et al. 2006). Even so, if we consider the hypothetically healthy population as a target for screening miR-519a-3p levels, it is important to ensure that other NDD that, like AD, course with self-aggregation proteins with spreading properties do not also converge in the upregulation of this miRNA. In this sense, miRNA dysregulation has previously been reported in both tauopathies and synucleopathies (Pratico 2020; Nies, Mohamad Najib et al. 2021). We have therefore considered it important to compare the miR-519a-3p levels found in some other non-AD tauopathies (CBD, GGT and PiD) and in some cases of synucleopathies (PD and PDD). To date, the presence of miR-519a-3p has never been analyzed in tauopathies, while in the case of synucleopathies, only analysis on IPSC-derived neurons from PD patients has been performed, and miR-519 reduction was reported (Tolosa, Botta-Orfila et al. 2018). Thus, our results point to increased miR-519a-3p levels as a specific sign of AD that serve as a promising tool when attempting to diagnose whether a person is in the asymptomatic stage of the disease. The next step is to start checking blood-derived samples of different cohorts that have been followed up longitudinally to implement the clinical use of miR-519a-3p in the diagnosis of AD.

## Conclusions

The present study points to miR-519a-3p as one of the elements responsible for PrP^C^ decrease in AD progression. The early expression of this miRNA in the disease could be of great interest in obtaining a specific asymptomatic AD biomarker that would allow for new fields of research aimed at discovering possible treatments.

## Acknowledgments

The authors thank Tom Yohannan for editorial advice and Miriam Segura-Feliu and Juan José López Jiménez for technical help. We also thank the Core facilities of IBEC for technical help.

## Credit authorship contribution statement: *Dayaneth Jácome*

Investigation, Methodology, Formal analysis. ***Tiziana Cotrufo:*** Investigation, Writing – review & editing. ***Pol Andrés-Benito:*** Investigation, Writing – review & editing. ***Eulàlia Martí:*** Software, Writing – review & editing ***Isidre Ferrer:*** Data curation, Writing – review & editing ***José Antonio del Río:*** Funding acquisition, Writing – review & editing ***Rosalina Gavín:*** Investigation, Formal analysis, Conceptualization, Writing - original draft, Writing - review & editing.

## Data availability statement

The data that support the findings of this study are available on request from the corresponding author, RG.

## Ethics approval statement

All postmortem human brains, from Clinic-IDIBAPS, HUB-ICO-IDIBELL, and NAVARRABIOMED Biobanks, were obtained following the Code of Ethics of the World Medical Association and the protocols of the local ethical committees.

## Abbreviations

Aβ,: β-amyloid;
AD,: Alzheimer’s disease;
CBD,: corticobasal degeneration;
CNS,: central nervous system;
CSF,: cerebrospinal fluid;
GGT,: glial globular tauopathy;
MCI,: mild cognitive impairment;
MMSE,: Mini-Mental Scale examination;
NDD,: neurodegenerative disease;
NFT,: neurofilament tangles;
nND,: non-neurodegenerative controls;
PD,: Parkinson’s disease;
PDD,: PD associated dementia;
PET,: positron emission tomography;
PHF,: paired helical filaments;
PiD,: Pick’s disease (PiD);
PrP^C^,: cellular prion protein.

## Supplementary material

**Supplementary Figure 1.**
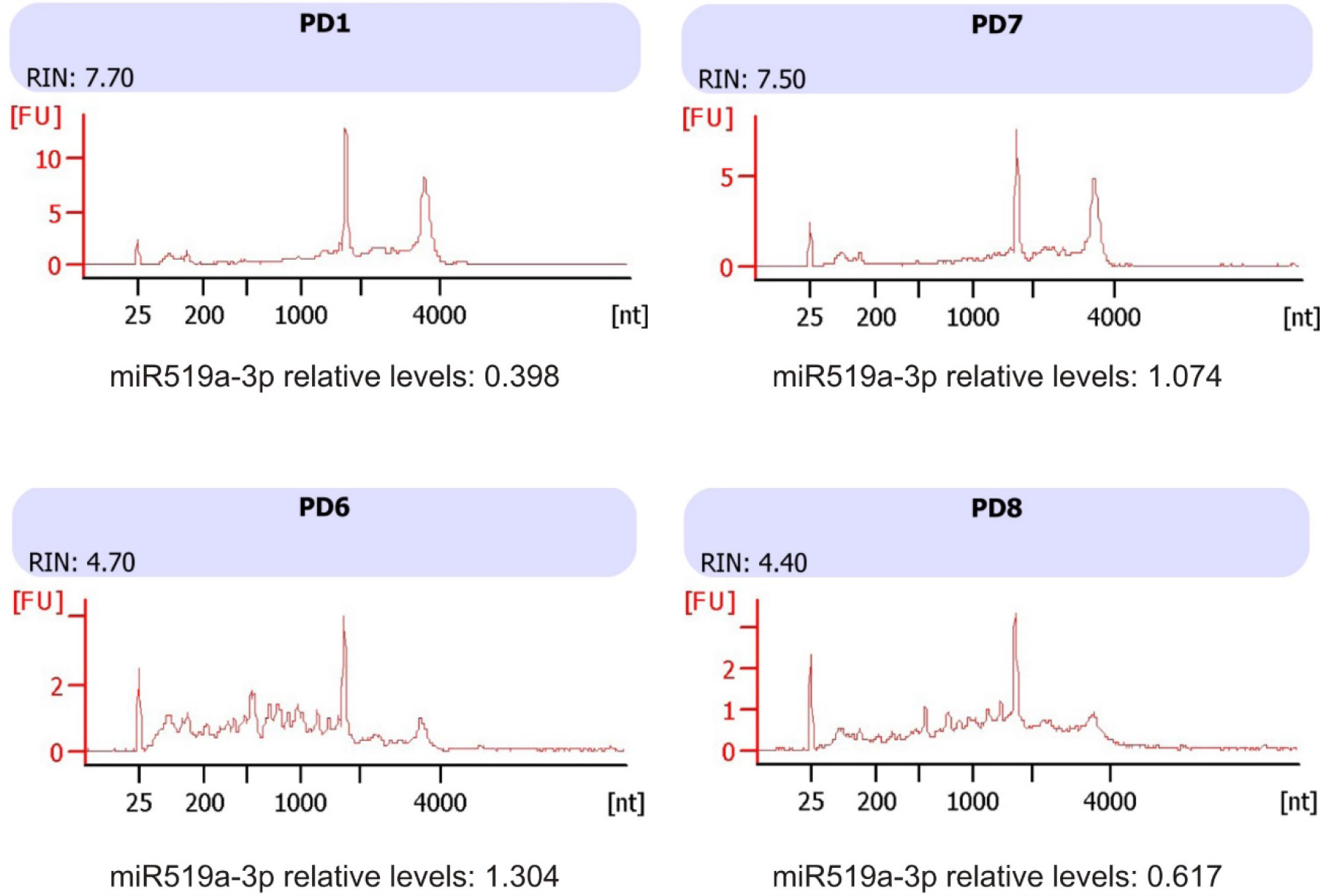
Results of total RNA analysis performed on 4 samples used in this study as example of non-association between RIN and miR-519a-3p levels.

**Supplementary Table 1.** Control (nND) and ND cortex samples from patients used in this study.

| Case number | Neuropathological Diagnosis | Gender | Age | Post-mortem delay | Biobank |
| --- | --- | --- | --- | --- | --- |
| C1 | nND | M | 59 | N.A. | HUB-ICO-IDIBELL |
| C2 | nND | M | 39 | 09h 15min | HUB-ICO-IDIBELL |
| C3 | nND | M | 62 | N.A. | HUB-ICO-IDIBELL |
| C4 | nND | F | 70 | N.A. | Clinic-IDIBAPS |
| C5 | nND | F | 81 | 23h 30min | Clinic-IDIBAPS |
| C6 | nND | F | 74 | 6h | Clinic-IDIBAPS |
| C7 | nND | M | 58 | 5h | Clinic-IDIBAPS |
| C8 | nND | M | 76 | 11h 30min | Clinic-IDIBAPS |
| C9 | nND | F | 74 | 00h 35min | Clinic-IDIBAPS |
| C10 | nND | M | 72 | N.A. | NAVARRABIOMED |
| C11 | nND | M | 79 | N.A. | NAVARRABIOMED |
| C12 | nND | M | 62 | N.A. | NAVARRABIOMED |
| C13 | nND | M | 91 | N.A. | NAVARRABIOMED |
| C14 | nND | F | 88 | N.A. | NAVARRABIOMED |
| AD1 | ADI A | F | 74 | 02h 45min | HUB-ICO-IDIBELL |
| AD2 | ADI A | F | 57 | 05h 00min | HUB-ICO-IDIBELL |
| AD3 | ADI 0 | M | 56 | 07h 10min | HUB-ICO-IDIBELL |
| AD4 | ADI A | M | 66 | 09h 45min | HUB-ICO-IDIBELL |
| AD5 | ADI A | M | 61 | 05h 35min | HUB-ICO-IDIBELL |
| AD6 | ADI 0 | F | 73 | 15h 45min | HUB-ICO-IDIBELL |
| AD7 | ADII 0 | F | 88 | 08h 00min | HUB-ICO-IDIBELL |
| AD8 | ADII 0 | M | 67 | 07h 15min | HUB-ICO-IDIBELL |
| AD9 | ADII A | M | 66 | 04h 55min | HUB-ICO-IDIBELL |
| AD10 | ADII A | F | 86 | 02h 15min | HUB-ICO-IDIBELL |
| AD11 | ADII 0 | M | 57 | 04h 30min | HUB-ICO-IDIBELL |
| AD12 | ADII A | M | 69 | 03h 45min | HUB-ICO-IDIBELL |
| AD13 | ADIII A | F | 82 | 02h 00min | HUB-ICO-IDIBELL |
| AD14 | ADIII 0 | M | 81 | 01h 30min | HUB-ICO-IDIBELL |
| AD15 | ADIII 0 | M | 66 | 05h 45min | HUB-ICO-IDIBELL |
| AD16 | ADIII A | F | 77 | 03h 10min | HUB-ICO-IDIBELL |
| AD17 | ADIII A | M | 64 | 06h 00min | HUB-ICO-IDIBELL |
| AD18 | ADIII0 | F | 79 | 03h 35min | HUB-ICO-IDIBELL |
| AD19 | ADIII B | F | 76 | 03h 50min | HUB-ICO-IDIBELL |
| AD20 | ADIII 0 | M | 83 | 07h 25min | HUB-ICO-IDIBELL |
| AD21 | ADIV A | F | 80 | 02h 45min | HUB-ICO-IDIBELL |
| AD22 | ADIV B | M | 84 | 12h 45min | HUB-ICO-IDIBELL |
| AD23 | ADIV C | F | 81 | 05h 00min | HUB-ICO-IDIBELL |
| AD24 | ADV C | F | 75 | 04h 15min | HUB-ICO-IDIBELL |
| AD25 | ADV C | M | 87 | 07h 05 min | HUB-ICO-IDIBELL |
| AD26 | ADV C | M | 74 | 10h 00min | Clinic-IDIBAPS |
| AD27 | ADV C | F | 75 | 08h 30min | Clinic-IDIBAPS |
| AD28 | ADV C | F | 88 | 09h 00min | Clinic-IDIBAPS |
| AD29 | ADV C | M | 78 | 05h 30min | Clinic-IDIBAPS |
| AD30 | ADV C | F | 89 | 08h 50min | Clinic-IDIBAPS |
| AD31 | ADV C | M | 91 | 07h 00min | Clinic-IDIBAPS |
| AD32 | ADV C | M | 92 | 07h 45min | Clinic-IDIBAPS |
| AD33 | ADV C | F | 87 | 03h 00min | Clinic-IDIBAPS |
| AD34 | ADV C | F | 78 | 09h 00min | Clinic-IDIBAPS |
| AD35 | ADV C | F | 87 | 00h 30min | Clinic-IDIBAPS |
| AD36 | ADV C | M | 90 | 05h 45min | Clinic-IDIBAPS |
| AD37 | ADV C | M | 73 | 06h 30min | Clinic-IDIBAPS |
| AD38 | ADV C | M | 89 | 08h 00min | Clinic-IDIBAPS |
| AD39 | ADV C | M | 79 | 04h 15min | Clinic-IDIBAPS |
| AD40 | ADV B | M | 78 | 09h 00min | Clinic-IDIBAPS |
| AD41 | ADV C | M | 83 | 06h 30min | Clinic-IDIBAPS |
| AD42 | ADV C | M | 88 | 03h 00min | Clinic-IDIBAPS |
| AD43 | ADV C | F | 90 | 05h 30min | Clinic-IDIBAPS |
| AD44 | ADVI C | F | 56 | 07h 00min | HUB-ICO-IDIBELL |
| AD45 | ADVI C | F | 71 | 05h 00min | Clinic-IDIBAPS |
| AD46 | ADVI C | F | 77 | 06h 15min | Clinic-IDIBAPS |
| AD47 | ADVI C | F | 90 | 00h 27min | Clinic-IDIBAPS |
| AD48 | ADVI C | F | 86 | 00h 37min | Clinic-IDIBAPS |
| AD49 | ADVI C | M | 81 | 00h 31min | Clinic-IDIBAPS |
| AD50 | ADVI C | F | 77 | 04h 30min | Clinic-IDIBAPS |
| AD51 | ADVI C | F | 82 | 07h 00min | Clinic-IDIBAPS |
| AD52 | ADVI C | F | 84 | 00h 32min | Clinic-IDIBAPS |
| AD53 | ADVI C | M | 82 | 05h 0min | Clinic-IDIBAPS |
| AD54 | ADVI C | M | 75 | 08h 15min | Clinic-IDIBAPS |
| AD55 | ADVI C | F | 95 | 05h 00min | Clinic-IDIBAPS |
| AD56 | ADVI C | F | 84 | 00h 37min | Clinic-IDIBAPS |
| AD57 | ADVI C | M | 85 | 04h 30min | Clinic-IDIBAPS |
| AD58 | ADVI C | F | 80 | 00h 23min | Clinic-IDIBAPS |
| AD59 | ADVI C | M | 73 | 05h 00min | Clinic-IDIBAPS |
| AD60 | ADVI C | F | 77 | 06h 00min | Clinic-IDIBAPS |
| AD61 | ADVI C | F | 76 | 00h 42min | Clinic-IDIBAPS |
| TAU1 | CBD | F | 69 | 03h 15min | NAVARRABIOMED |
| TAU2 | CBD | M | 85 | 04h 00min | NAVARRABIOMED |
| TAU3 | CBD | M | 55 | 10h 00min | NAVARRABIOMED |
| TAU4 | GGT | M | 47 | 13h 20min | NAVARRABIOMED |
| TAU5 | GGT | F | 61 | 05h 20min | NAVARRABIOMED |
| TAU6 | GGT | F | 85 | 05h 00min | NAVARRABIOMED |
| TAU7 | PiD | F | 77 | 02h 20min | NAVARRABIOMED |
| TAU8 | PiD | F | 80 | 08h 00min | NAVARRABIOMED |
| TAU9 | PiD | F | 65 | 04h 00min | NAVARRABIOMED |
| TAU10 | PiD | F | 80 | 04h 35min | NAVARRABIOMED |
| PD1 | PD3-4 | M | 68 | 09h 20min | Clinic-IDIBAPS |
| PD2 | PD3-4 | F | 70 | 04h 30min | Clinic-IDIBAPS |
| PD3 | PD4 | F | 69 | 04h 30min | Clinic-IDIBAPS |
| PD4 | PD4 | F | 84 | 04h 30min | Clinic-IDIBAPS |
| PD5 | PD4-5 | F | 81 | 08h 15min | Clinic-IDIBAPS |
| PD6 | PD5 | M | 81 | 05h 30min | Clinic-IDIBAPS |
| PD7 | PD5 | F | 70 | 08h 30min | Clinic-IDIBAPS |
| PD8 | PD5 | M | 71 | 05h 30min | Clinic-IDIBAPS |
| PD9 | PD5 | F | 77 | 03h 30min | Clinic-IDIBAPS |
| PD10 | PDD5 | M | 72 | 07h 45min | Clinic-IDIBAPS |
| PD11 | PD5 | M | 83 | 14h 00min | Clinic-IDIBAPS |
| PD12 | PD5 | F | 88 | 07h 00min | Clinic-IDIBAPS |
| PD13 | PD5 | F | 77 | 07h 30min | Clinic-IDIBAPS |
| PD14 | PDD5 | M | 81 | 05h 00min | Clinic-IDIBAPS |
| PD15 | PD5-6 | F | 77 | 05h 45min | Clinic-IDIBAPS |
| PD16 | PDD6 | M | 80 | 07h 30min | Clinic-IDIBAPS |
| PD17 | PD | F | 79 | 02h 20min | NAVARRABIOMED |
| PD18 | PD | F | 78 | 03h 45min | NAVARRABIOMED |
| PD19 | PD | M | 84 | 07h 00min | NAVARRABIOMED |
| PD20 | PD | M | 90 | 08h 00min | NAVARRABIOMED |
| PD21 | PD5 | M | 87 | N.A. | NAVARRABIOMED |

*nND: Non-neurodegenerative; AD: Alzheimer's disease; CBD: Corticobasal degeneration; GGT: Glial globular tauopathy; PiD: Pick's disease; PD: Parkinson's disease; PDD: Parkinson's disease with dementia; F: female; M: male; N.A.: not available*

